# Cellular Responses to Photothermal Therapy: Heat-Induced ERK Signaling and Intercellular Communication in Solid Tumors

**DOI:** 10.1101/2025.02.27.640541

**Authors:** Farsai Taemaitree, Yuta Takano, Daisuke Yamaguchi, Kazushi Yamaguchi, Yudai Yamashita, Motosuke Tsutsumi, Chentao Wen, Kohei Otomo, Kenji Hirai, James Hutchison, Indra Van Zundert, Sandra Krzyzowska, Maria Bravo, Sayuki Hirano, Koutarou D Kimura, Kazuhiro Aoki, Susana Rocha, Tomomi Nemoto, Beatrice Fortuni, Hiroshi Uji-i

## Abstract

Nanoparticle-mediated photothermal therapy (PTT) shows promise as a standalone cancer treatment but faces clinical challenges due to inconsistent efficacy. Its translation is further hindered by a limited understanding of plasmon-induced heat effects, such as stress responses and intercellular signaling. Here, we investigate how plasmon-induced local heating affects cellular behavior and fate within tumor spheroids by focusing on the activity of the extracellular signal-regulated kinase (ERK). Spheroids were prepared from HeLa cells that express a FRET-based ERK sensor, and ERK activity changes under photothermal stimuli were tracked using a deep-learning program, 3DeeCellTracker. Gold nanostars were used as highly efficient photothermal transducers. Our results revealed significant alterations in ERK signaling patterns upon photothermal stimulation compared to spontaneous ERK activity in untreated spheroids, including changes in the activation frequency, timing, and duration. Notably, photothermal-induced ERK activity propagated across neighboring cells within the spheroid, suggesting intercellular communication. Furthermore, analysis of cell death and division further demonstrated that laser power modulates cellular fate during photothermal therapy. This study provides insights for predicting the therapeutic effects of PPT and guides the rational design of next-generation photothermal strategies. Additionally, our approach demonstrates the potential of FRET-based biosensors and deep-learning tools as powerful methods to study the effects of various therapeutic stimuli on solid tumors at the single-cell level.

## Introduction

Effective eradication of tumor cells is a critical goal in cancer therapy, with photothermal therapy (PTT) emerging as a particularly promising method. PTT uses photothermal agents to generate localized heat upon laser irradiation, where precise modulation of parameters – such as light intensity – determines whether a pathological state or cell death is induced.^1–3^ Surface plasmon-mediated PTT,^4^ which involves metal nanoparticles (NPs), is gaining increasing attention for its ability to drastically enhance thermal effects at the nanometer scale through strong light absorption and scattering in the visible and near-infrared range.^5^ Although the development of photothermal nanoagents is progressing rapidly, the key cellular and molecular mechanisms triggered by the plasmon-induced hyperthermia in tumors are still not characterized. Bridging this knowledge gap is crucial for advancing the next-generation of cancer treatments, as it requires a deeper insight of how heat stimuli affect cellular behavior.

One effective method for exploring cellular responses to stress is by examining extracellular signal-regulated kinase (ERK) activity. ERK belongs to the family of mitogen-activated protein kinases (MAPK), which play a crucial role in pathways that transmit extracellular signals to their intracellular targets,^6–9^ thereby regulating numerous biological processes including cell proliferation and differentiation, cellular adaptation/survival,^10^ and apoptosis.^11,12^ Therefore, monitoring the activation of ERK signaling pathways holds promise for gaining valuable insights into how cells adapt and respond to localized thermal stress at a single cell level.

Concurrently, the implementation of advanced three-dimensional (3D) tumor platforms represents an innovative approach to address this challenge, enhancing the relevance of preclinical studies. Among various 3D cancer models, spheroids effectively mimic the intricate architecture of cancerous tissues and their microenvironment, displaying cell-to-cell interactions, as well as intercellular signaling.^13–15^

This study proposes a novel technique to examine cellular responses to photothermal therapeutic stress by monitoring ERK activity in 3D tumor spheroids. Our unique approach enabled the comprehensive evaluation of plasmon-induced PTT effects in HeLa spheroids by integrating the use of our recently developed biosensor for ERK activity,^16^ with our advanced 3D image analysis tool, 3DeeCellTracker.^17^ Gold nanostars (AuNSs) were selected as photothermal transducers,^18^ and ERK activity was monitored using two-photon microscopy.^19^ This integrated approach offers several advantages: (i) allowing for precise control of laser intensity to regulate heat generation by AuNSs, (ii) facilitating high-resolution image acquisition in multicellular 3D cell systems, and (iii) enabling single-cell level analysis of over 1000 cells simultaneously. The results obtained elucidate how cells respond to nanoparticle-induced PTT in solid tumors, providing crucial information for next-generation treatments, and setting a new standard for investigating complex biological processes in 3D models.

## Methods

### General

Absorption spectra were recorded using a UV-Vis spectrophotometer (V-730, JASCO, Japan). Dynamic Light Scattering (DLS) and zeta potential measurements were conducted at room temperature in Milli-Q water using Zetasizer Nano ZS (Malvern Panalytical Ltd., U.K.). All chemicals and solvents used in this study were of analytical grade and used as received unless otherwise stated.

### Synthesis and characterization of Cysteamine-coated AuNS (AuNS-Cys)

Synthesis: a gold nanostar (AuNS) colloidal solution was prepared according to previous reports.^18^ Briefly, 3 mL of 1.0% citric acid solution was added to 20 mL of boiling 1.0 mM tetrachloroauric(III) acid trihydrate (HAuCl_4_) aqueous solution and stirred for 15 min at 110°C. The dispersion was allowed to cool to room temperature and was subsequently stored at 4°C for 24 h. Next, 10 mL of 0.25 mM HAuCl_4_ aqueous solution, 100 μL of 1.0 M HCl, and 100 μL of the gold colloid dispersion were mixed and stirred. 100 μL of 3.0 mM silver nitrate (AgNO_3_) aqueous solution and 100 μL of 100 mM ascorbic acid (AA) solution were added simultaneously to the mixture at room temperature. Immediately after the mixture turned blue, it was centrifuged at 3000 × g for 15 min to stop nucleation, then the supernatant was removed. Milli-Q water was added and a dispersed AuNS solution was obtained after sonication for 5 min.

The AuNS dispersion (1.0 mg/mL) was mixed with 80 mM cysteamine aqueous solution in equal volumes, sonicated for 30 min under light shielding, centrifuged at 3000 × g for 10 min, and washed with Milli-Q water. This process was repeated three times to remove excess cysteamine and obtain cysteamine-coated AuNS (AuNS-Cys) dispersed in Milli-Q water at 1.0 mg/mL concentration.

#### STEM characterization

10 μL of AuNS-Cys dispersion was dropped onto a copper mesh grid (collodion film-applied mesh, Nissin EM Co., Ltd.), dried under vacuum for at least 30 min, and observed using a HITACHI HD-2000 (Hitachi, Japan) with an acceleration voltage of 200 kV. Images were acquired in multiple fields of view for one grid and used to measure particle size. Several fields of view were measured and the particle diameter was determined as the maximum Euclidean distance between any two points on the AuNS-Cys perimeter. The particle core diameter was the darkest area from the shading of the STEM image. The results showed that the mean particle diameter ± standard deviation (SD) was 101.3 ± 15.0 nm (n = 131) and the mean core diameter was 37.8 ± 6.5 nm (n = 131).

### Evaluation of local heating effects

8 uL of the AuNS-Cys dispersion was dropped onto a cover glass and spin-coated at 30 rps for 30 s. The samples were then dried on a hotplate at 90°C for 30 s. A polydimethylsiloxane (PDMS) framework was pasted onto the cover glass to function as a chamber for poly (N-isopropylacrylamide) (PNIPAM) solution. Next, 40 µL of a solution of water and 1,4-dioxane (1:1 v/v) containing 5 mg/mL PNIPAM (Sigma-Aldrich, Japan) was added to the PDMS framework, covering the spin-coated AuNS-Cys. Plasmonic heating effects were observed using an inverted optical microscope (Ti-U, NIKON, Japan) with an 808 nm CW laser, 12.5 mW. The laser beam was reflected by a dichroic mirror and focused onto the sample using an objective lens (PlanFluo x60, NA 0.85, Nikon). To eliminate Rayleigh scattering, a long-pass filter (ET750SP-2P8, Chroma, USA) was placed in front of the camera when using the 850 nm fs laser. Additionally, a long-pass filter (ET500LP, Chroma) was placed in front of the camera to cut off the light of the 850 nm harmonic.

Bright-field images from the microscope illumination were analyzed by software (Hokawo/Hamamatsu Photonics, Japan) using a CCD camera (ImagEM, Hamamatsu Photonics). Laser power measurements were performed at the objective lens using a power meter (THORLABS, USA) to determine the power of the laser incident on the sample.

### Culturing HeLa cells with a stable expression of EKAREV-NLS (#T-HeLa)

For the real-time observation of ERK activity in individual cells, a fluorescence resonance energy transfer (FRET) sensor protein, EKAREV-NLS, was stably expressed in HeLa cells (#T-HeLa).^20^ Briefly, plasmids with EKAREV-NLS construct developed previously^16^ and plasmid with PiggyBac transposon were introduced into cells via lipofection (Lipofectamine, Thermo Fischer Scientific, USA) at 3 to 1 ratio. After lipofection, the cells were cultured in the presence of 10 μg/mL blasticidin (Invivo Gen, USA) to select transfected cells. Cells stably expressing EKAREV-NLS were maintained and passaged in Dulbecco’s modified Eagle’s minimal essential medium (DMEM, Wako, Japan) supplemented with 10% fetal bovine serum (FBS, Gibco, USA) and 1% penicillin/streptavidin (P/S, Wako, Japan). Cells were cultured at 37°C in 5% CO_2_/air. Cell passages were performed at 80-90% confluency using trypsin.

### Dark cytotoxicity assay

The cytotoxicity of AuNS-Cys was evaluated by measuring the number of viable cells with a cell counter. In detail, the cells were seeded in 35 mm glass bottom dishes at a density of 2.0 × 10^5^ cells per dish with 1 mL cell culture medium and incubated at 37°C for 24 h under 5% CO_2_. The next day, after confirming cell adhesion to the bottom of the dish, 50 μL of AuNS-Cys solution (1.0 mg/mL) was added and incubated for 10 min. After incubation, the cells were washed with Dulbecco’s Phosphate-Buffered Saline (D-PBS (−), Wako, Japan) and incubated for 48 h at 37°C. Next, the number of viable cells was counted by staining dead cells with trypan blue. Each test was performed in triplicate.

### Preparation of spheroids of #T-HeLa

A suspension of #T-HeLa cell culture medium (200 μL) containing 3600 cells was dropped into a 96-well plate designated for spheroid preparation (MS-9096U; Sumitomo Bakelite, Japan). The plates were then covered with lids and incubated for 48 h at 37°C in 5% CO_2_. Spheroid formation was monitored using an optical microscope.

### 3D-imaging of the spheroids of #T-HeLa by a two-photon laser microscope

AuNS-Cys solution was added to each well of a 96-well plate (MS-9096U) filled with #T-HeLa spheroids and gently pipetted, followed by incubation for 10 min at 37°C and 5% CO_2_. The spheroids were removed from the wells and submerged in 3 mL of D-PBS (−) solution to remove excess AuNS-Cys. The spheroids were then transferred to a 35 mm glass bottom dish coated with poly-lysine, and 2.0 mL of culture medium was added. Typically, the dose concentration of the AuNS-Cys dispersion solution was 25 μg/mL for the experiments at high concentration of AuNS-Cys and 2.5 ng/mL for those at low concentration of AuNS-Cys.

The sample was then imaged using a multi-photon excitation inverted microscope (A1R-MP, Nikon, Japan) equipped with a silicone oil immersion objective lens (CFI Plan Apochromat Lambda S 25XC SilNA 1.05, SIL, Nikon, Japan). Correction rings were aligned to maximize the fluorescence intensity of the samples. A wavelength-tunable Titanium Sapphire (Ti-Sa) laser (850 nm, 120 fs, 80 MHz) was used to excite cyan fluorescent protein (CFP), yellow fluorescent protein (YFP), and AuNS-Cys. The excitation laser powers used were: 1.5 mW mm^-2^ at the objective for most cases (defined as normal conditions), 75 mW mm^-2^ under intense conditions, and 25 mW mm^-2^ under moderate laser power conditions. Note that the excitation wavelength of 850 nm can simultaneously realize photo-thermal local stimulation by AuNS-Cys and imaging of ERK activity through two-photon excitation. Dichroic mirrors were used to detect the fluorescence and photoluminescence (PL) from the FRET probe and AuNS-Cys, respectively. 400-492 nm fluorescence was acquired for CH1 (CFP), 500-550 nm for CH2 (YFP), and 601-657 nm for CH3 (AuNS-Cys, although detectable in all three channels).

The acquired area was 512 × 512 pixels (related to 450 × 450 μm^2^) in the xy plane. In the z-direction, imaging was performed from the cover glass surface (the bottom of the spheroid) to the center, which is typically approximately 100 μm in height. During measurements, a stage incubator was used to maintain the extracellular environment at 37°C and 5% CO_2_. The cells were placed on the stage for 30 min prior to imaging for stabilization.

### Image analysis of the spheroids of #T-HeLa

Changes in ERK activity were visualized as ratiometric images obtained using ImageJ-Fĳi.^21^ First, background signals from the CFP and YFP were eliminated by automatic thresholding. For each image, the fluorescence intensity of the YFP channel was divided by that of the CFP channel, obtaining ratiometric images representing the FRET efficiency. A color bar was added to each image, indicating the magnitude of FRET efficiency, which is in turn associable with the ERK activity.

The 3DeeCellTracker^17,22^ was employed to extract changes in FRET efficiency for each cell within a cancer spheroid. A deep neural network, 3D U-Net^23^, was trained using one or a few 3D raw images along with their corresponding annotations of the cell regions. The trained network was then used to detect and segment the cells in a series of 3D images. Another neural network, FFN^17^, was used to track the dynamic positions of cells. From detection to tracking, the process demonstrated an accuracy of 97.6 ± 1.9% (n = 3) in the present cases.

The data on changes in FRET efficiency from the cell nuclei were further processed to extract information on the timing of ERK activity. Initially, a linear de-trending process was applied to the FRET efficiency data from individual cell nuclei to remove any linear trends. The mean values and standard deviations (SD) of FRET efficiency were then calculated for each cell. FRET efficiencies that exceeded the mean of the control + 2SD were considered a significant ERK activity, and information on the timing and frequency of such activity was then extracted.

## Results and discussion

### Characterization of Gold Nanostars and their photothermal properties

Gold nanostars (AuNSs) were chosen as photothermal transducers because of their characteristic plasmonic band in the near-infrared (NIR) light region (∼850 nm) and their excellent photothermal conversion efficiency (PCE).^18^ They also exhibit an intrinsic photoluminescence (PL) under two-photon excitation^24^, which enables their direct observation with fluorescent microscopy.

AuNSs were synthesized via seed-mediated growth, as previously reported.^18^ To improve stability in aqueous media, enhance interaction with the cell membrane, and increase cellular uptake, AuNSs were coated with cysteamine, resulting in positively charged nanoparticles at neutral pH.^25^ The star-shaped morphology of AuNS-Cys was confirmed by Scanning Transmission Electron Microscopy (STEM) (Figure 1B) and their average diameter was estimated to be ∼100 nm (Figure S1). UV-Vis spectroscopy revealed a typical extinction spectrum of star-shaped NPs, characterized by a very broad plasmonic band localized in the NIR region, with a peak at 890 nm (Figure 1C).^26^ The change in surface charge after cysteamine functionalization was determined by zeta potential measurements. The obtained values for AuNS and AuNS-Cys in deionized water were –40.4 ± 4.9 mV and +19.5 ± 5.1 mV, respectively (Figure 1D), confirming the positive shift in surface charge upon functionalization.

**Figure 1.**
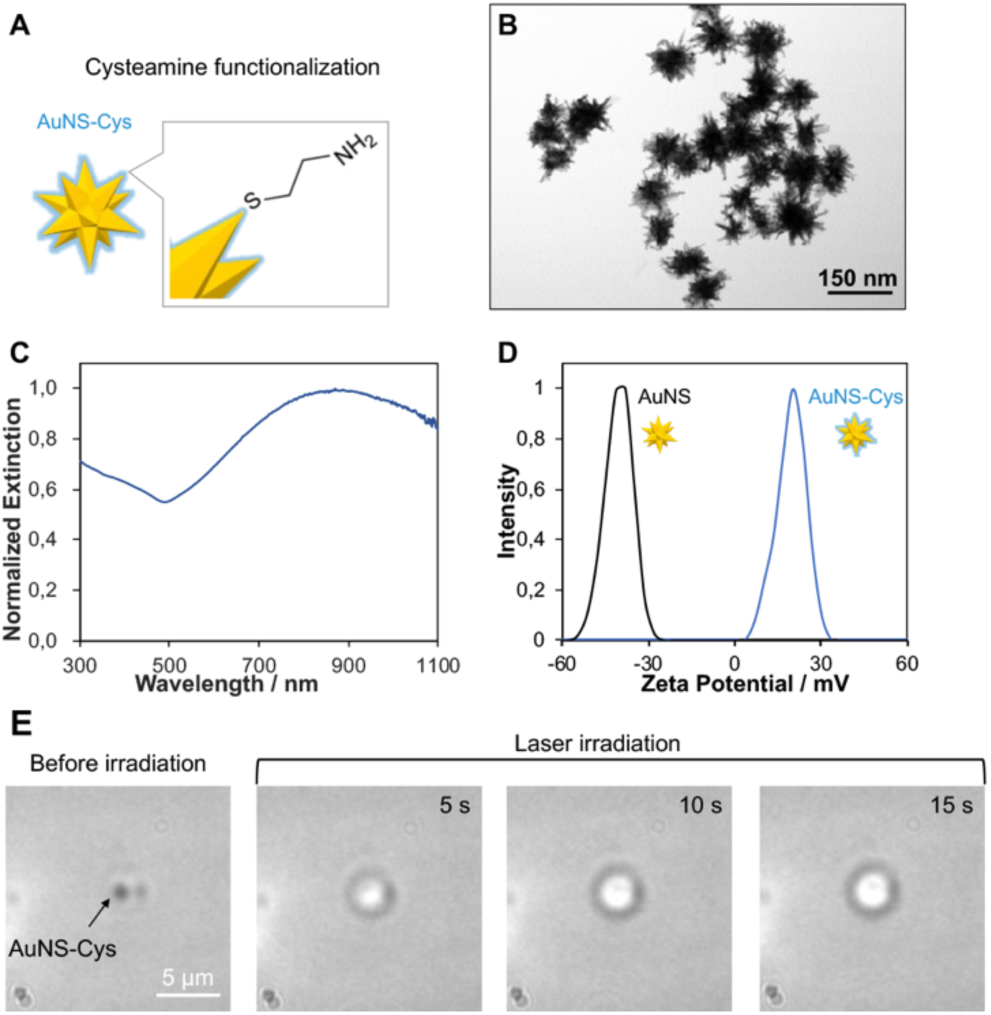
Characterization of AuNS-Cys. (A) Schematic representation of AuNS functionalization with cysteamine. (B) STEM image of AuNS-Cys. (C) Extinction spectrum of AuNS-Cys in water. (D) Zeta-potential profiles of AuNS (black) and AuNS-Cys (blue). (E) Thermally-induced phase separation in PNIPAM aqueous solution upon photothermal conversion of AuNS-Cys.

The local heating effect of AuNS-Cys was verified by inducing the phase separation in the aqueous solution of a thermoresponsive polymer, poly(N-isopropylacrylamide) (PNIPAM). PNIPAM is known to exhibit a lower critical solution temperature (LCST) of approximately 45°C when dissolved in a 1:1 (v/v) mixture of water and dioxane.^27^ To demonstrate the photothermal effect, AuNS-Cys were mixed with a 5 mg/mL PNIPAM solution in 1:1 (v/v) water:dioxane and irradiated with 808 nm excitation. After 5 s of irradiation focused on AuNS-Cys, phase separation was observed, appearing as bubble formation around the AuNSs (Figure 1E). The phase separation continued to enlarge over 15 s of irradiation, indicating that plasmon-induced heating from the AuNS raised the temperature of their surroundings to at least 45°C within a few seconds of NIR irradiation. The efficiency of AuNS photothermal conversion was then calculated by previously reported methods^28^ and estimated to be 33.7% (details of the calculation are reported in the SI and Figure S2). This value outperformed most recently reported photo-thermal materials.^29–30^ The phase separation of PNIPAM aqueous solution and the high estimated conversion efficiency indicate that AuNSs can efficiently generate heat upon NIR excitation and can therefore be applied for studying the cellular response to heat in 3D cancer models.

### Monitoring ERK activity in 3D cancer spheroids using FRET sensor proteins

In order to explore the cellular response to PTT in tumors, real-time monitoring of ERK activity was performed in spheroids upon plasmonic-induced local heat by AuNS-Cys. ERK activation was tracked by encoding a previously reported FRET-based biosensor for ERK activity (EKAREV-NLS)^16^ in HeLa cells prior to forming spheroids. The EKAREV-NLS sensor includes a FRET donor-acceptor pair (a cyan-emitting protein, CFP, and a yellow-emitting protein, YFP), a sensory domain in between the two fluorescent proteins which is susceptible to phosphorylation, and a ligand domain separated by a flexible linker. Upon perceiving stress, the sensory domain is phosphorylated and binds to its ligand domain. This interaction results in a conformational change, bringing the two fluorescent proteins closer together, thereby increasing the rate of FRET events (Figure 2A).^16^ By observing the change in FRET efficiency as intensity ratio between the acceptor and the donor (YFP/CFP), the ERK activity can be estimated.^31^Click or tap here to enter text. The modified cells are designated as #T-HeLa cells in this study and were used to form 3D tumor spheroids in a low attachment 96-well plate.

**Figure 2.**
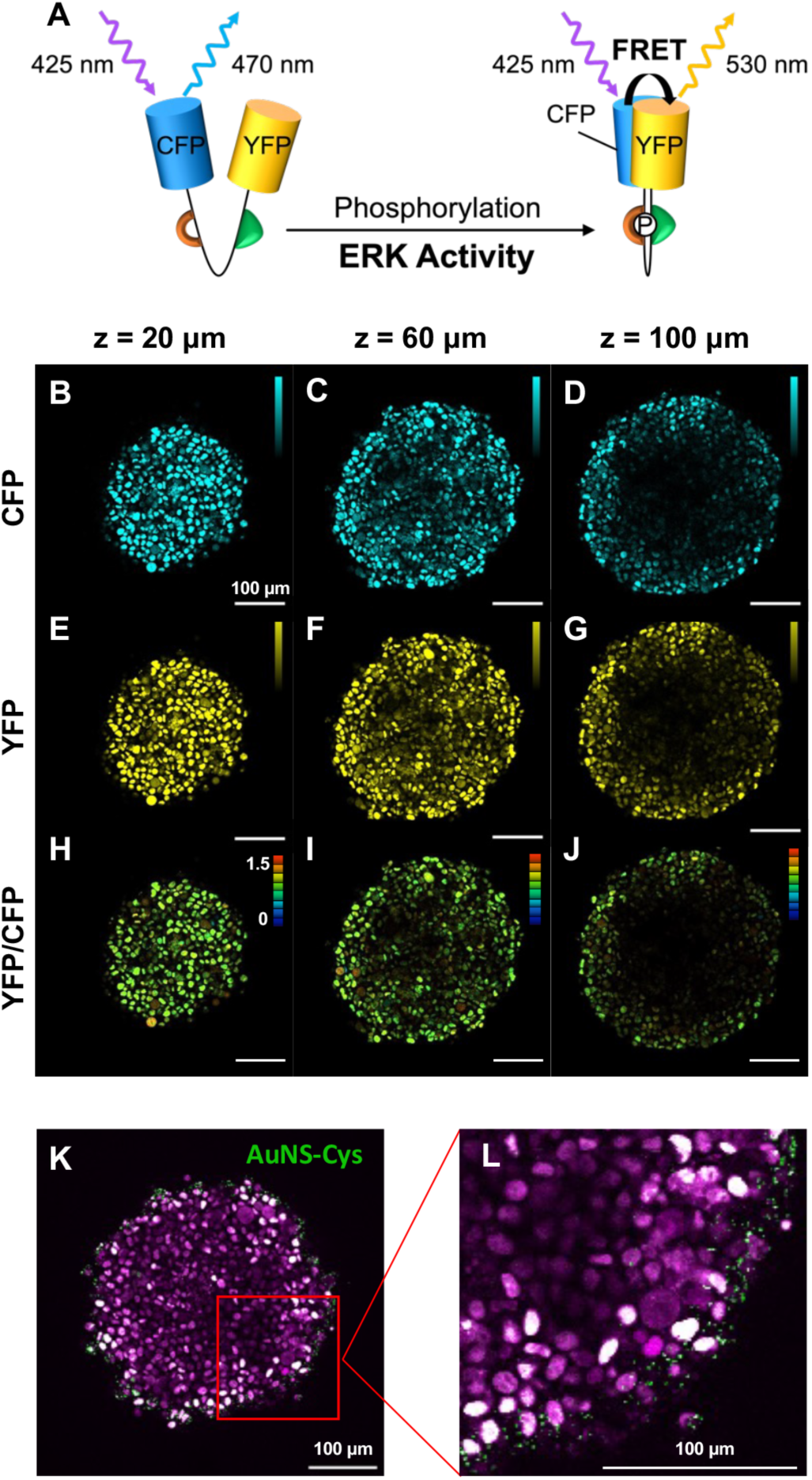
Schematic representation of the FRET biosensor (EKAREV-NLS) structure and activation mechanism. (A). xy fluorescence images extracted from z-stacks of EKAREV-NLS expressing HeLa (#T-HeLa) spheroids at 20, 60 and 100 μm depth from the spheroid surface displaying CFP emission (B-D, respectively) and YFP emission (E-G, respectively), upon two-photon donor excitation at 850 nm. YFP/CFP ratiometric images indicating the FRET efficiency corresponding to the ERK activity (H-J) (color bar YFP/CFP: 0-1.5). Low and high-magnification xy fluorescence images extracted from a z-stack of a #T-HeLa spheroid after 10 min of incubation with AuNS-Cys upon two-photon excitation at 850 nm (K and L, respectively): cells observed in magenta (YFP emission), whereas the AuNS-Cys photoluminescence is displayed in green (CFP emission was also detected in the green channel with the CFP and YFP overlap appearing in white).

To verify the successful expression of the ERK FRET sensor in #T-HeLa cells and to evaluate the influence of two-photon imaging on #T-HeLa, steady-state ERK activity in #T-HeLa spheroids was monitored using two-photon fluorescence microscopy. The fluorescence signals of both fluorescent proteins of the sensor, CFP and YFP, were clearly detected when the donor was excited (Figure 2B-D and E-G, respectively), indicating the successful expression of the sensor and the occurrence of FRET. The use of two-photon excitation enabled the observation of sharp fluorescence signals at 20 and 60 μm depth from the spheroid surface (Figure 2B,E and C,F, respectively). At 100 μm depth, the signal was slightly lower in intensity at the spheroid core but still clearly observable (Figure 2D,G). The FRET efficiency variations, corresponding to different levels of ERK activity, were analyzed using YFP/CFP ratiometric imaging (Figure 2H-J). In the ratiometric images, a high YFP/CFP ratio, with cells appearing orange-to-red, indicates high ERK activity, whereas a low YFP/CFP ratio, with cells observed in green, reflects low ERK activity. Figure 2H-J shows that in a NP-free spheroid, during the scanning, most cells exhibited low ERK activity, appearing in green. Interestingly, a small ratio of cells within the spheroid was observed in orange-red (Figure 2H-J), indicative of ERK activation. These results align with recent studies reporting that spontaneous activation or propagation from neighboring cells can result in sporadic pulses of ERK activity.^7,32^ These experiments confirm that the proposed method enables the visualization of FRET efficiencies in individual cells within a spheroid, thereby allowing for precise monitoring of the ERK signaling activity in a 3D tumor structure at the single-cell level.

To accumulate AuNS-Cys into #T-HeLa spheroids, AuNS-Cys were added to the spheroid-containing medium at a final concentration of 25 μg/mL and incubated for 10 min. Note that dark cytotoxicity assays (Figure S3) confirmed that AuNS-Cys (25 μg/mL) had no significant impact on cell viability or ERK activity after 48 h without laser irradiation. Prior to fluorescence imaging, the samples were washed to remove excess NPs. On the fluorescence microscope, AuNS-Cys were visualized by their intrinsic PL upon 850 nm two photon excitation,^18^ whereas cells were observed though CFP and YFP emission. Confocal fluorescence images revealed that, after 10 min incubation, AuNS-Cys were primarily localized in the peripheral layer of the spheroid, with some adhering to the cell membrane and others being internalized (Figure 2K-L). The efficient interaction between AuNS-Cys and the cells was attributed to the cysteamine coating. The positive surface charge of AuNS-Cys, acquired through cysteamine functionalization, facilitates electrostatic interactions with the negatively charged cell membrane^33^, thereby enhancing NP uptake.^34^ This data indicates that AuNS-Cys were successfully accumulated in #T-HeLa spheroids and can therefore be used for studying the effect of plasmonic-induced heat on ERK activity in spheroids over time.

To monitor the spontaneous ERK activity in #T-HeLa spheroids over time without nanoparticles (control), fluorescence scans of a selected focal plane at 100 μm from the spheroid surface were performed at intervals of 2 min for 2 h. Ratiometric fluorescence images at 0, 2 and 10 min are reported as exhibiting the most significant changes (Figure 3 A-C, respectively). According to the data shown in Figure 2, the majority of cells were inactive over the 10-min interval, appearing in green, while a small proportion of active cells were observed in orange or red. This activity pattern remained stable over time, with the same cells exhibiting high ERK activity throughout the 10-min period.

**Figure 3.**
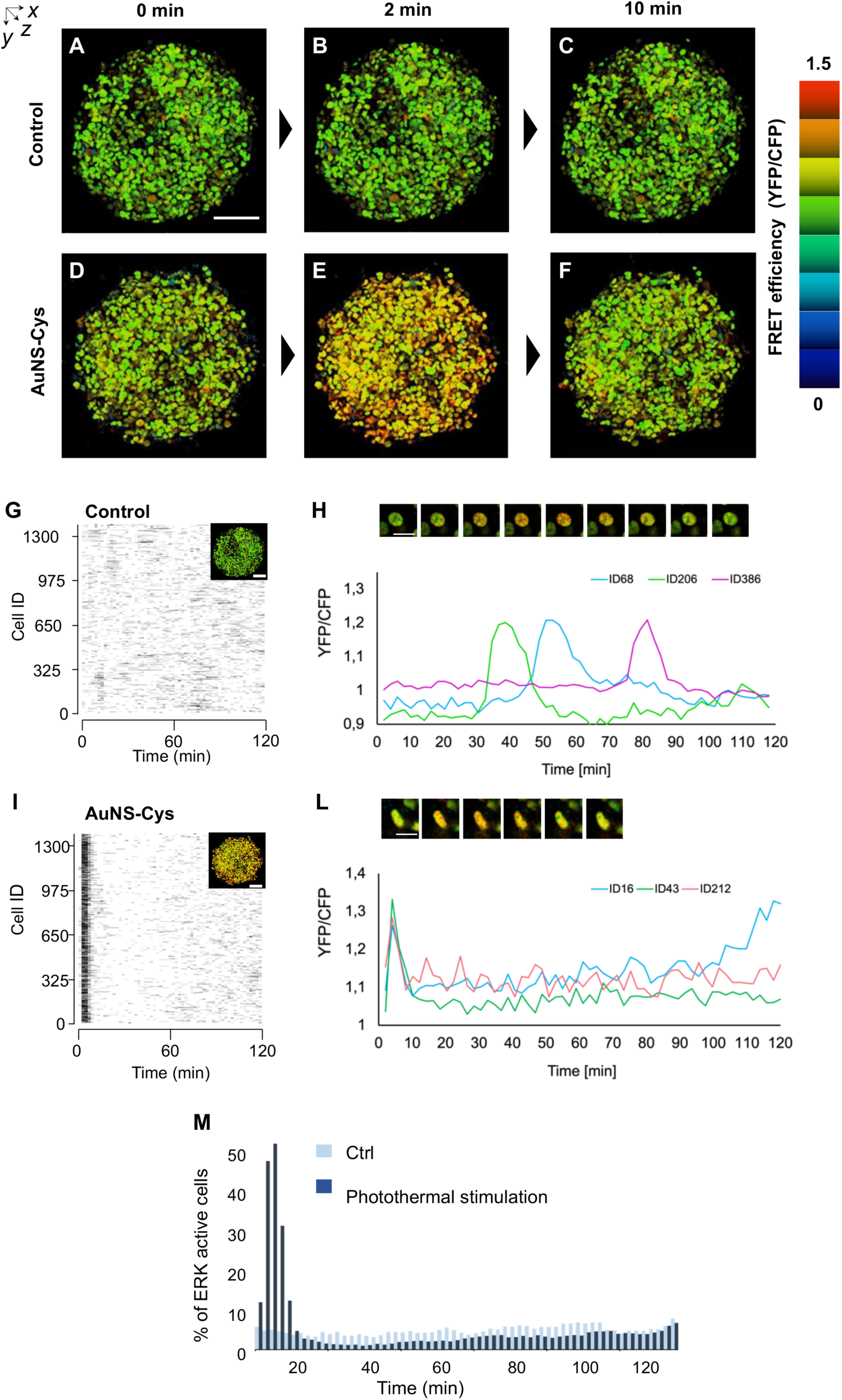
Two-photon fluorescence microscopy. Ratiometric maximum intensity z-projections of #T-HeLa spheroids as control (A-C), and of #T-HeLa spheroids incubated with AuNS-Cys (D-F) at 0 min (first scan, A and D, respectively), after 2 min from the first scan (B and E, respectively) and after 10 min from the first scan (C and F, respectively) with 850 nm two photon excitation (scale bar: 100 μm). 3DeeCellTracker analysis. Raster plots showing the timing of ERK activation in single cells as a function of time in the control (G) and upon AuNS-Cys-induced photothermal stimuli (I). The black horizontal lines indicate the ERK-activated state, determined by the FRET efficiency of YFP/CFP exceeding the mean of the control +2 SD, and the duration of each back line is 2 min. FRET efficiency as YFP/CFP over time of 3 individual cells selected within the spheroids reported in A-C for the control (H) and D-F for the AuNS-Cys-induced heat condition (L). Insets show ratiometric images of a single representative cell per condition captured at 2-min intervals (scale bar: 20 μm). Percentage of ERK-active cells as a function of time (M) for the control (dark blue) and for AuNS-Cys-induced heat condition (cyan), obtained from G and I, respectively.

In order to evaluate the effect of plasmon-induced heat on the spontaneous ERK activity observed in the control, the same experiment was repeated after incubation with AuNS-Cys. Ratiometric fluorescence images showed that right after AuNS-Cys irradiation (Figure 3D), most cells, which are initially in a low ERK activity state, displayed some activation, appearing yellow. This is an apparent difference from the control, which is overall more green (Figure 3A). The results suggest that the immediate heat generation may have initiated ERK activation shortly after irradiation. The plasmon-induced heat effect became more pronounced 2 min after the initial irradiation (Figure 3E), evidenced by a significant increase in ERK-active cells, reflected by the orange-to-red shift in FRET efficiency. Notably, by 10 min post-irradiation, most cells reverted to their initial ERK-inactive state (Figure 3F), with FRET efficiency levels in green, comparable to those of the control (Figure 3C). This dynamic response highlights the transient nature of the ERK activation induced by plasmonic heating.

In this study, the use of AuNS-Cys as thermal transducers enabled the delivery of a single, non-reproducible thermal stimulus. Upon irradiation with fs-pulsed laser light, AuNS-Cys likely underwent morphological changes,^35^ causing a shift in their plasmonic band and reducing their photothermal conversion efficiency. This was evidenced by a significant decrease in PL intensity after the initial scans (Figure S4) and confirmed by the PNIPAM phase separation assay, which could not be repeated using the same AuNS-Cys after the first irradiation. These observations indicate that plasmon-induced heating was a one-time event.

These data demonstrate that plasmon-induced heating elicits rapid and robust activation of ERK in tumor spheroids, which is transient, as most cells revert to an ERK-inactive state within 10 min.

### 3DeeCellTracker analysis of ERK activity in 3D spheroids at single cell-level

To evaluate the timing and duration of ERK activation at the single-cell level within spheroids, fluorescence imaging data were analyzed using 3DeeCellTracker, an automatic deep-learning program developed by our research group. This tool integrates 3D U-Net^23^ and FFN^17^ networks trained on relevant datasets, enabling highly accurate (97.6 ± 1.9%) cell nucleus detection and tracking with minimal parameter adjustments. Cell nuclei were identified from YFP fluorescence using 3D U-Net + watershed^36^, and their positions were predicted over time for tracking. The fluorescence intensities of CFP and YFP were then extracted from each tracked cell to calculate the FRET efficiency (YFP/CFP). Using 3DeeCellTracker, over 1300 cells were simultaneously tracked for 120 min, providing robust insights into ERK activation dynamics. The resulting data was visualized as raster plots, illustrating the temporal dynamics of the ERK activation in individual cells within the spheroid, for both control (Figure 3G) and AuNS-Cys-induced heat (Figure 3I). Cells with high FRET efficiencies (exceeding the mean of the control + 2 SD) were classified as ERK-active and represented as black lines in the plots, where each line corresponds to a 2-min duration. These raster plots corroborate the patterns observed in the fluorescence images: the control exhibited random ERK activation events, with black lines appearing sporadically across different cells over the 120-min interval, while the AuNS-Cys-induced heat triggered a rapid, synchronized ERK activation in the majority of cells, evidenced by densely aligned black lines within the first few minutes post-irradiation. To further characterize the dynamics of ERK activation, FRET efficiency as YFP/CFP over time of three representative cells from control spheroid and spheroid with AuNS-Cys was analyzed (Figures 3H and L, respectively). In the control, spontaneous ERK pulses appeared as broader peaks with durations of 20.0 ± 2.0 min, distributed randomly over the 120-min interval (Figure 3H). In contrast, heat-induced ERK activation displayed sharp peaks of shorter duration (9.3 ± 3.1 min) that were consistently localized within the first few minutes following AuNS-Cys irradiation (Figure 3L). These findings demonstrate that plasmon-induced photothermal effects rapidly cause synchronized ERK activation in the majority of cells and reveal that heat-stimulated ERK activity is significantly shorter in duration than the spontaneous ERK activity in unperturbed cells. This unprecedented insight is essential for understanding and predicting heat-induced cellular responses in solid tumors, as the timing and duration of ERK activation are critical determinants of key cellular processes, including proliferation, differentiation, and apoptosis.^11,37^

The percentage of ERK-active cells in the control and following plasmon-induced heat was calculated and compared over time (Figure 3M). The basal ERK activity was detected in 3.9 ± 0.6% of cells in the control group. This low activity is comparable to that observed in 2D cell monolayers nearing confluency, where ERK activity diminishes as cells experience contact inhibition of growth – a regulatory mechanism that controls proliferation.^7^

In contrast, the transient increase in ERK activity observed within a few minutes after AuNS-Cys irradiation occurred in 53.7 ± 11.0% of the spheroid cells. Interestingly, 15 min after the initial activation, the cellular ERK activity significantly dropped, resulting in a percentage of activation lower than that of the control. After 95 min, ERK activity returned to levels comparable to the control, indicating a transient suspension of ERK activity following plasmon-induced heat. Two mechanisms could explain this phenomenon: (1) temporary cell inactivation caused by photothermal stimuli, as protein denaturation and cell inactivation are known to occur when tissue temperatures reach 42°C,^38^ or (2) negative feedback suppressing ERK activity in the Ras/ERK or MEK/ERK pathways triggered by local photothermal stimulation.^39,40^ Although further investigations are required to confirm these hypotheses, this study provides unique direct observations of ERK activity dynamics in response to local heat in a spheroid over time.

### ERK signaling and intercellular communication

To investigate the intercellular communication underlying ERK activation in response to local photothermal stimuli, the experiments were repeated using a 10 times lower concentration of AuNS-Cys (2.5 ng/mL). This adjustment ensured that only a few cells within the spheroid interacted with AuNS-Cys, allowing for the analysis of ERK activation in neighboring cells that were not in direct contact with AuNS-Cys (Figure 4). Fluorescence ratiometric images at the first scan (0 min) showed a uniformly low ERK activity state (with cells in green) and confirmed that only a few AuNS-Cys accumulated in the spheroid, as indicated by white arrows in Figure 4A. Two min after the initial scan, only cells in the surrounding area of AuNS-Cys (white squares, Figure 4B) showed remarkable ERK activation, appearing orange-red, while in the rest of the spheroid, the ERK activation level remained stable (in green).

**Figure 4.**
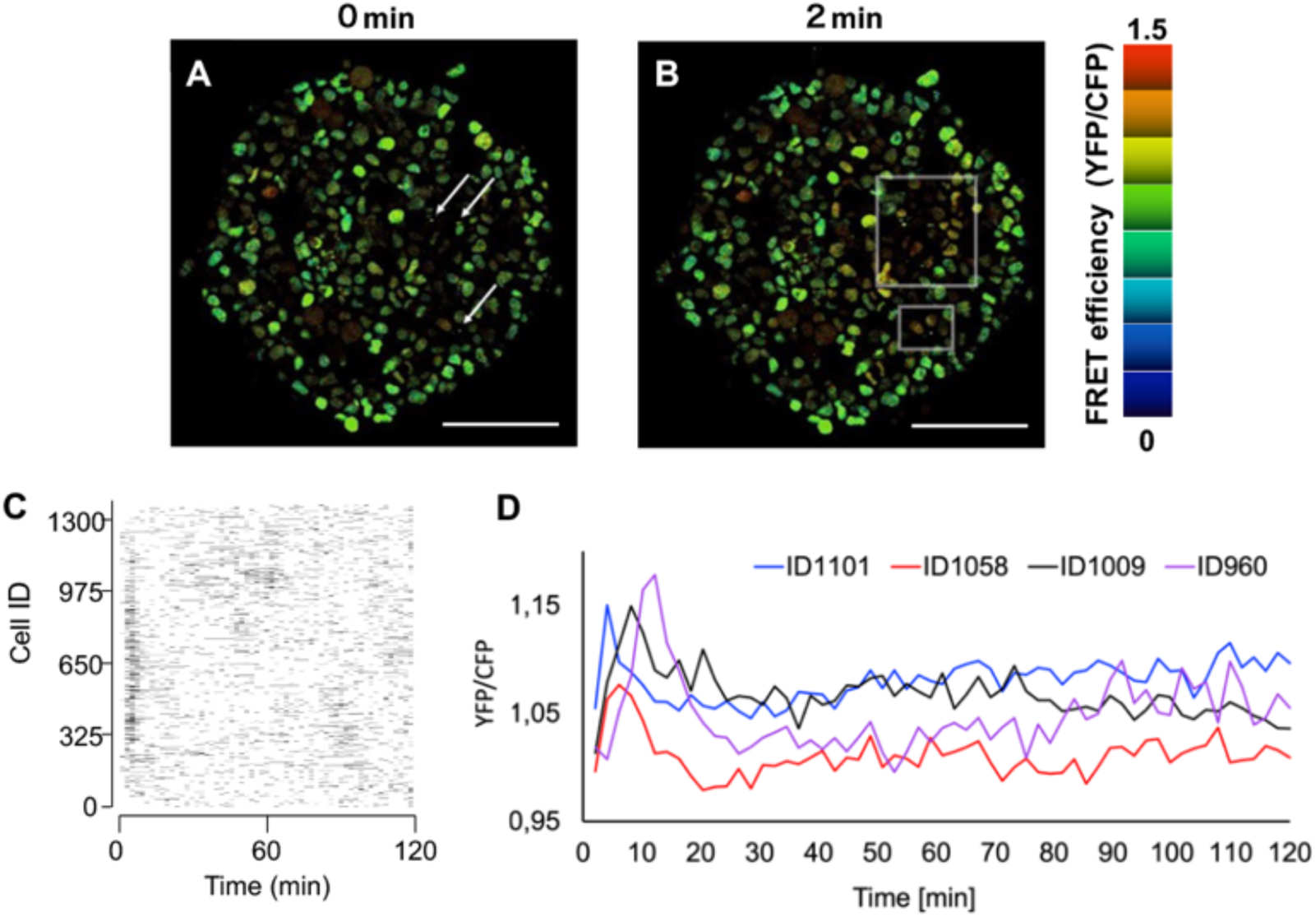
Ratiometric xy fluorescence images extracted from a z-stack of a #T-HeLa spheroid incubated with a reduced concentration of AuNS-Cys at 0 min. (first scan, A) and 2 min after the initial scan (B), with white arrows indicating the AuNS-Cys location and white squares delimiting the heat-induced ERK activation area. Corresponding raster plot obtained from 3DeeCellTracker analysis showing the timing of ERK activation in single cells as a function of time (C). The black horizontal lines indicate the ERK-activated state, determined by the FRET efficiency of YFP/CFP exceeding the mean of the control +2 SD, and the duration of each back line is 2 min. FRET efficiency as YFP/CFP over time of three individual cells selected within the white squared area of the spheroid (D).

The raster plot obtained from 3DeeCellTracker (Figure 4C), although very similar to that observed at higher AuNS-Cys concentrations, exhibits a less pronounced activation within the first few minutes following the initial scan because of the lower concentration of AuNS-Cys used. The variation of the YFP/CFP ratio in three individual cells over time (Figure 4D), selected from the area surrounding AuNS-Cys (white squares, Figure 4B), revealed distinct sharp activation peaks occurring at different time points within the 15-min interval (maximum ratio values at 2, 8, and 12 min). These variations indicate a discrepancy in the timing of ERK activation among neighboring cells. Given that heat propagates through biological tissue at a rate of tens of micrometers per millisecond,^41,42^ the observed delay in ERK activation among adjacent cells cannot be attributed to heat propagation. Instead, this delay likely arises from a stimulus-induced ERK signaling cascade mediated by intercellular communication. Previous studies have shown that ERK signaling can propagate between cells via mechanisms by which epidermal growth factor receptor (EGFR) ligands cleaved through ERK-induced ADAM17 activate EGFR in neighboring cells.^7,32,43^ These mechanism can result in coordinated responses across a cell population. The delayed ERK activation in the neighboring cells, observed here, highlights the complexity of ERK activation, which can be triggered by both direct heat effects and secondary signaling through cell-to-cell communication, potentially influencing cell fate. Similar patterns have been observed in studies in 2D cell monolayer where mechanical or chemical stimuli triggered wave-like ERK activation spreading across tissues.^7^

### Linking ERK activity to cellular fate

The investigation of ERK activity in a 3D cell system under plasmon-induced heating provided valuable insights into ERK signaling dynamics. However, the therapeutic potential of AuNS-Cys as a photothermal agent remains to be evaluated. To this end, spheroids were incubated with a high concentration of AuNS-Cys (25 µg/mL) to ensure extensive coverage. Sulforhodamine B (SRB), a red-fluorescent marker that selectively accumulates in cells with severely compromised membranes,^44^ was added to the cell culture medium as an indicator for cell death upon photothermal stimulation.

To determine the extent of photothermally induced cell death evoked by our previously set experimental conditions used for the heat-induced ERK activity experiments (Figure 3), control spheroids and spheroids containing AuNS-Cys were exposed to laser irradiation at 1.5 mW mm⁻² and imaged at 0 min, 2 min, and 10 min (Figure 5A–C and D–F, respectively). At the initial scan, a baseline level of dead cells, appearing in red, was observed in both conditions and remained stable over the 10-min interval. This is an expected phenomenon in unperturbed spheroids, where a certain proportion of cell death naturally occurs due to limitations in nutrient and oxygen diffusion across the densely packed 3D cellular structure, leading to apoptotic and necrotic processes.^45^ At this laser power, no significant increase in cell death was observed in the presence of AuNS-Cys, except for a single cell undergoing death after 2 min (marked with a light blue circle in Figure 5D–E). Applying an additional 3 s of irradiation at 25 mW mm⁻² laser power prior to imaging (at 1.5 mW mm⁻²) did not produce an immediate effect at 0 min (Figure 5G). However, after 2 min, a subset of cells underwent cell death (marked with light blue circles in Figure 5G–H), with a progressive increase over time, as additional cells exhibited cell death by 10 min (marked with green circles in Figure 5H–I). This increase in cytotoxicity can be attributed to the photothermal effect of AuNS-Cys. The cytotoxic response was even more pronounced when spheroids containing AuNS-Cys were irradiated at 75 mW mm⁻² for 3 s prior to imaging. Under these conditions, a substantial percentage of cells exhibited cell death, as indicated by extensive red fluorescence within the spheroid structure (Figure 5J–L). The proportion of dead cells increased linearly over time, with the majority of the cells in the spheroid appearing red by 10 min post-scan (Figure 5L), demonstrating a significant photothermal-induced cytotoxic effect at this laser power.

**Figure 5.**
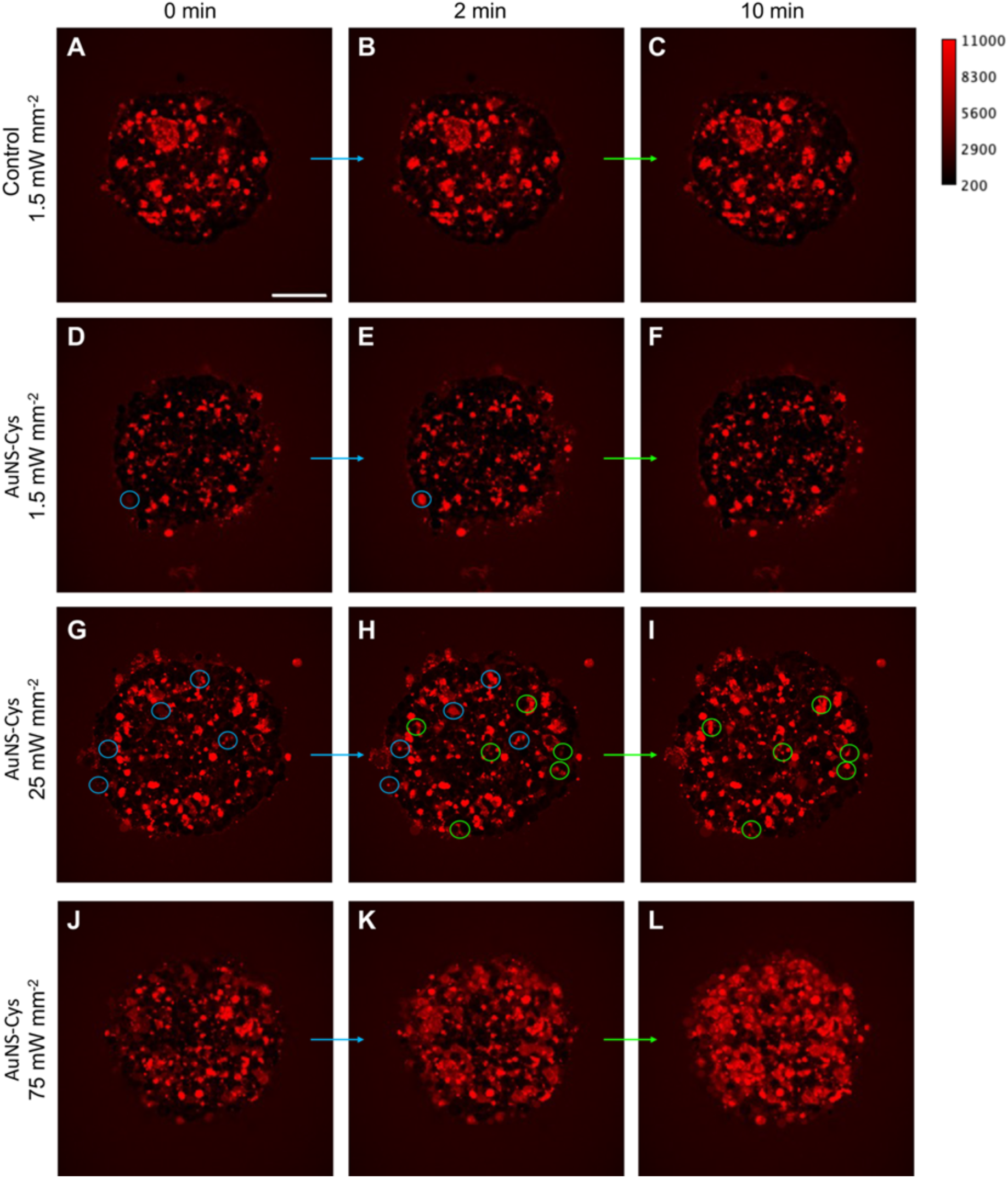
Spheroid cell death assay using sulforhodamine B (SRB) staining. Confocal xy fluorescence images extracted from z-stacks show #T-Hela spheroids under different conditions: control spheroids without AuNS-Cys, imaged after the first scan at 1.5 mW mm⁻² (0 min) and at 2 min and 10 min post-scan (A–C, respectively); spheroids with AuNS-Cys, imaged after the first scan at 1.5 mW mm⁻² (0 min) and at 2 min and 10 min post-scan (D–F, respectively); spheroids with AuNS-Cys pre-irradiated for 3 s at 25 mW mm⁻², then imaged under the same condition at 0 min, 2 min, and 10 min (G–I, respectively); spheroids with AuNS-Cys pre-irradiated for 3 s at 75 mW mm⁻², then imaged under the same condition at 0 min, 2 min, and 10 min (J-L, respectively). Scale bar: 100 μm. Two-photon excitation: 850 nm.

To investigate the correlation between photothermal-induced ERK activity and cell death, FRET efficiency (YFP/CFP) in #T-HeLa cells was assessed following an additional 3-s irradiation at 25 mW mm⁻² prior to imaging. This laser power induced moderate levels of cell death (Figure 5G-I), whereas higher power settings (75 mW mm⁻²) were not feasible due to excessive cell mortality (Figure 5J-L), which would have compromised fluorescence imaging. At 25 mW mm⁻², ratiometric imaging revealed a high proportion of ERK-active cells (Figure S5), with raster plot analysis indicating that 72.2% of cells initiated ERK activity within two min from the initial scan. This activation rate was significantly higher than that observed under standard photoexcitation conditions (53.7% at 1.5 mW mm⁻²), where no significant cell death was detected (Figure 5D-F). These findings suggest that cell death within spheroids may be associated with an increased proportion of ERK-active cells, corroborating the link between ERK signaling and photothermal-induced cell death.

Furthermore, the rate of cell division under ERK experiment conditions (1.5 mW mm⁻² laser power) was assessed over 2 h post-irradiation. Interestingly, a significant increase in cell proliferation was observed compared to steady-state conditions (control), with 102.7 ± 6.4 cell divisions recorded among 1300 cells within this period (Figure S6). This percentage was substantially higher than that of the control (74.7 ± 9.9), suggesting that at this laser power, plasmon-induced heating promotes cell division rather than cell death. Given that ERK signaling is implicated in cell proliferation and survival pathways^10,37^, these results confirm the pivotal role of ERK activity in determining cellular fate, where laser power modulation dictates the balance between photothermal-induced death and proliferation. Such insights are crucial for refining NP-mediated PTT to achieve controlled therapeutic effects, while minimizing unintended cellular damage.

## Conclusions

This study presents a groundbreaking approach for monitoring photothermal effects on cellular behavior within solid tumors by tracking variations in ERK signaling. By employing gold nanostars as thermal transducers, a FRET biosensor to visualize ERK activity in a 3D cell model, and the advanced deep-learning tool 3DeeCellTracker for data analysis, we have uncovered unprecedented insights into how cells within a spheroid respond to localized heat.

The key findings were that (i) ERK activity occurs spontaneously and randomly in a limited number of cells within unperturbed spheroids; (ii) plasmon-induced heat drastically increases the number of ERK-activated cells; (iii) heat-induced ERK activation is synchronized among spheroid cells and occurs within a few minutes from the stimulation; (iv) heat-triggered ERK activation is shorter in duration than spontaneous activation; (v) after heat stimuli cells undergo transient ERK inactivation; (vi) ERK signals propagate several micrometers from the heat source through intercellular communication; and (vii) the laser power used for photothermal stimulation critically determines cellular fate, influencing either proliferation or cell death.

These findings offer critical insights into the spatiotemporal dynamics of ERK signaling under photothermal conditions, providing a foundation for the rational design and optimization of next-generation cancer photothermal therapies. By advancing our understanding on how cellular behavior is modulated by heat at the molecular and multicellular levels, this study paves the way for more precise and effective therapeutic strategies against solid tumors.

## Acknowledgments: Funding

We acknowledge financial support from MEXT under the JSPS KAKENHI (21H01753 and 21K19036 to Y.T., 22K20524 to F.T., 22K14578, 19K15406 to M.T., JP22KK0100 to M.T., K.O., K.K., 22H02756 to K.O., 20H05669 to K.O. and T.N., 16H06545 to K.D.K., JP22H04926, 22H02625 to K.A., 21H04634, 23H04877, 23K17856 to H.U., 23H04877 to K.H. and H.U.), Bilateral Program (JPJSBP120232301 to Y.T and S.R.), Core-to-Core Program for Advanced Research Network (CCA20190003), and the Cooperative Research Program of “NJRC Mater. & Dev.”, JST CREST (JPMJCR20E4 to K.O.), a grant of Joint Research by the National Institutes of Natural Sciences (01112002), Grant-in-Aid for Research at Nagoya City University (48, 1912011, 1921102), Joint Research of the Exploratory Research Center on Life and Living Systems (ExCELLS) (22EXC201, 23EXC204, 24EXC201), and Dr. H. Mitomo and Ms. C. Takeuchi for the technical support for DLS and zeta potential measurements. We acknowledges financial support from Research Foundation of Flanders (FWO) research grants (G0D4519N, G081916N, VS08523N, G0C1821N, G022724N), postdoctoral fellowship (12X1419N, 12X1423N to B.F., 12A6N25N to I.V.Z.), and from the KU Leuven (C14/15/053, C14/19/079, C14/22/085, C14/23/090).). F.T. acknowledges financial support from Nakatani Foundation. We thank the Open Facility, Global Facility Center, Creative Research Institution, Hokkaido University for allowing us to use SEM and STEM. The authors would like to thank the NIKON Imaging Center at Hokkaido University for the imaging equipment and software.

## Author Contributions

H.U., B.F., and T.N. conceived the idea and directed the project. F.T., Y.T., D.Y., Y.Y., M.T., and K.H. synthesized the compounds and acquired and analyzed the data. Y.T., F.T., D.Y., K.Y., M.T., K.O., M.B., S.K., I.V.Z., B.F., and S.R. performed microscopy experiments. Y.T., F.T., D.Y., B.F., and H.U. wrote the manuscript draft. F.T., D.Y., K.Y., M.T., C.W., K.O., K.H., J.H., K.D.K, K.A., and T.N. developed the analysis methods using the FRET sensor and deep-learning program. All the authors contributed to editing the manuscript.

## Data availability

All data generated or analyzed during this study are included in this published article and its Supplementary Information file.

