## Supplementary material for "Cellular Responses to Photothermal Therapy: Heat-Induced ERK Signaling and Intercellular Communication in Solid Tumors": SI

Supporting information

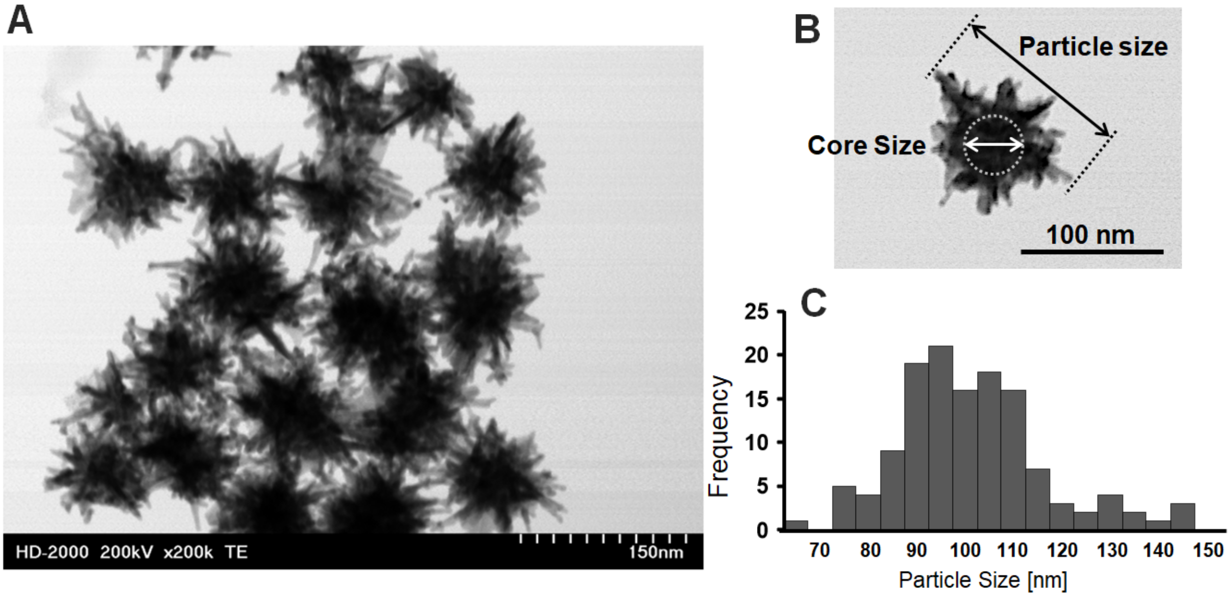


**Figure S1.** (A) STEM image of AuNS-Cys. (B) Schematic representation of the diameter calculation for the core and the particle: the particle diameter was estimated as the maximum Euclidean distance value between any two points on the AuNS-Cys perimeter. The particle core diameter was obtained from the darkest area from the shading of the STEM image. (C) Particle size distribution profile extracted from STEM images: mean particle diameter ± standard deviation was 101.3 ± 15.0 nm (n = 131) and the mean core diameter ± standard deviation was 37.8 ± 6.5 nm (n = 131).

Photothermal conversion efficiency (PCE)

The photo-thermal conversion (η) was determined according to the previously described method[28];

$$\eta = \frac{hS(T_{Max} - T_{Surr}) - Q_{Dis}}{I(1 - 10^{-A_{700}})} \qquad eq\ (1)$$

where h is the heat transfer coefficient, S is the surface area of the container, and $hS$ is determined from Equation (2). $T_{Max}$ is the maximum temperature achieved by illumination, and $T_{Surr}$ is the temperature of the surroundings. $Q_{Dis}$ represents the heat dissipated from the laser, mediated by the solvent and container. $I$ corresponds to the intensity of the power source (700 nm laser at 0.030 W), and $A_{700}$ represents the absorbance of the sample solution at 700 nm.

$$hs = \frac{m_D\, C_D}{\tau_s} \qquad eq(2)$$

$m_D$ and $C_D$ are the mass and heat capacity of deionized water, respectively, and $\tau_s$ is the sample system time constant, which is determined according to Equation (3) and the slope shown in Figure. S7.

$$t = -\tau_s\, In(\theta) \qquad eq(3)$$

θ is a dimensionless parameter, known as the driving force temperature, as calculated using equation (4)

$$\theta = \frac{T - T_{surr}}{T_{max} - T_{surr}} \qquad eq(4)$$

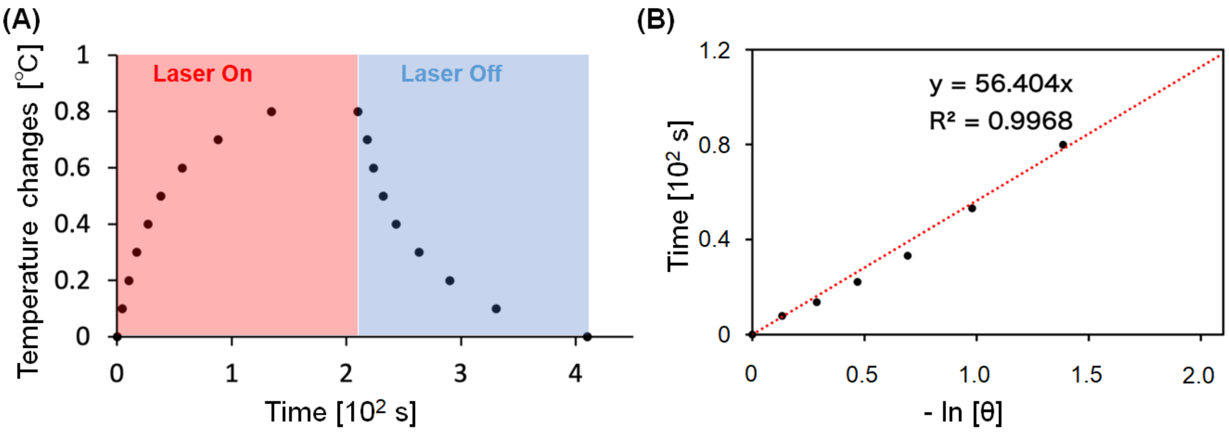


**Figure S2.** (A) Heating-cooling experiment under 30 mW 700 nm CW laser irradiation to determine the photothermal conversion efficiency (PCE). The laser was switched off after the sample reached a steady-state temperature of 220 s. (B) Linear fitting for the time and -ln θ plot. Linear time data versus -ln θ obtained from the cooling period in panel (A).

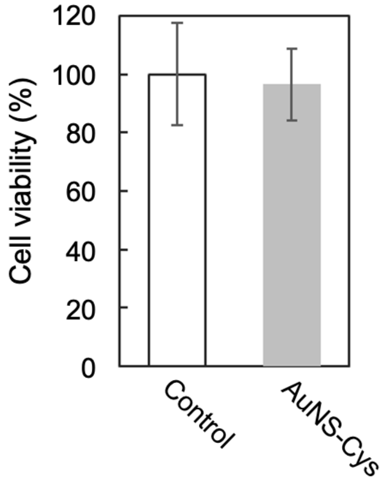


**Figure S3**. Dark cytotoxicity assay showing the viability of #T-HeLa cells with or without treatment with AuNS-Cys after 48 h incubation at 37°C.

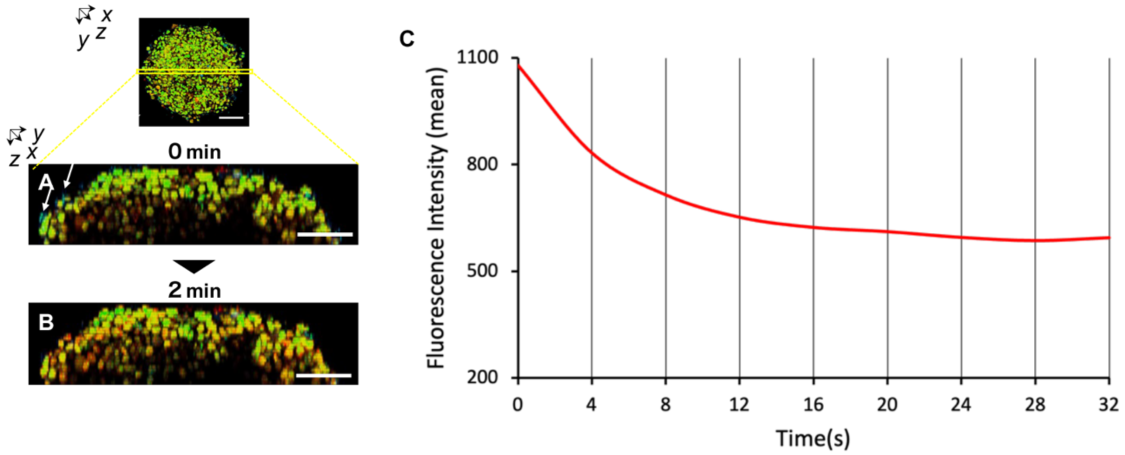


**Figure S4.** Maximum intensity z-projection (y-z) of the spheroids corresponding to Figure 3D,E (A and B, respectively), with the white arrows indicating the PL of AuNS-Cys. Photoluminescence intensity of AuNS-Cys over time upon irradiation with an 850 nm fs-pulsed laser, 22 mW mm$^{-2}$ (B).

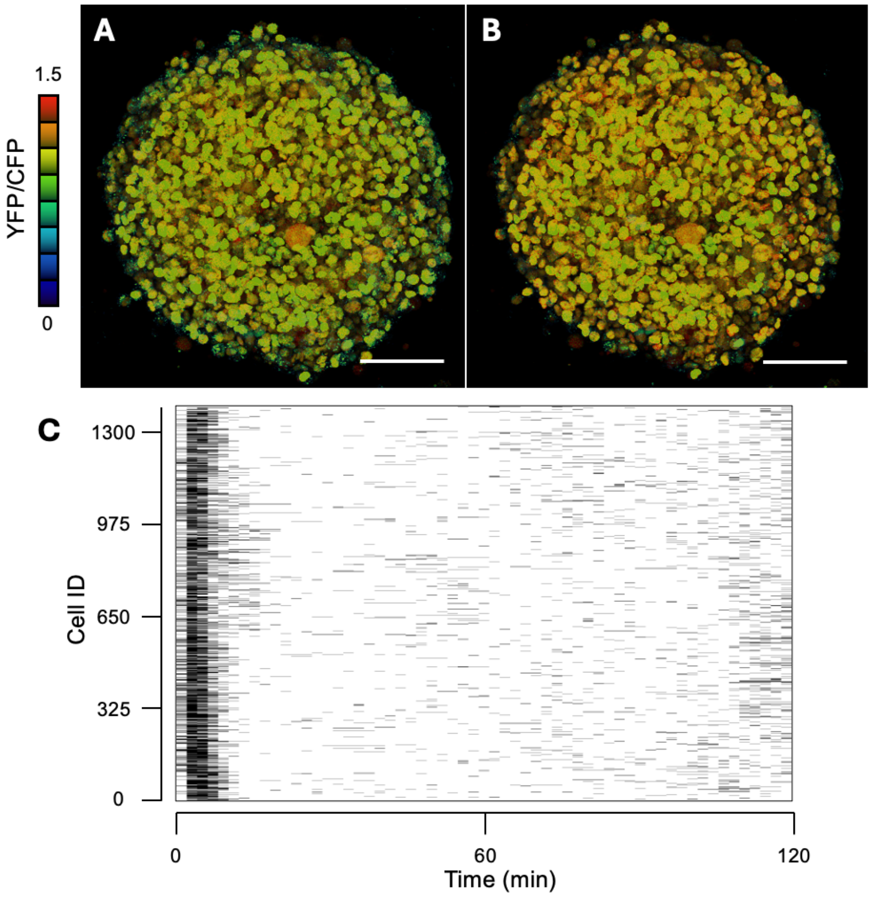


**Figure S5**. Ratiometric maximum intensity z-projections of #T-HeLa spheroids incubated with AuNS-Cys (two-photon excitation at 850 nm, 22 mW $mm^{-2}$) at (A) 0 min and (B) 2 min. (C) Raster plots obtained from the 3DeeCellTracker analysis showing the timing of ERK activation in single cells as a function of time. The black horizontal lines indicate the ERK-activated state, determined by the FRET efficiency of YFP/CFP exceeding the mean of the control +2 SD, and the duration of each black line is 2 min. Scale bar = 100 μm.

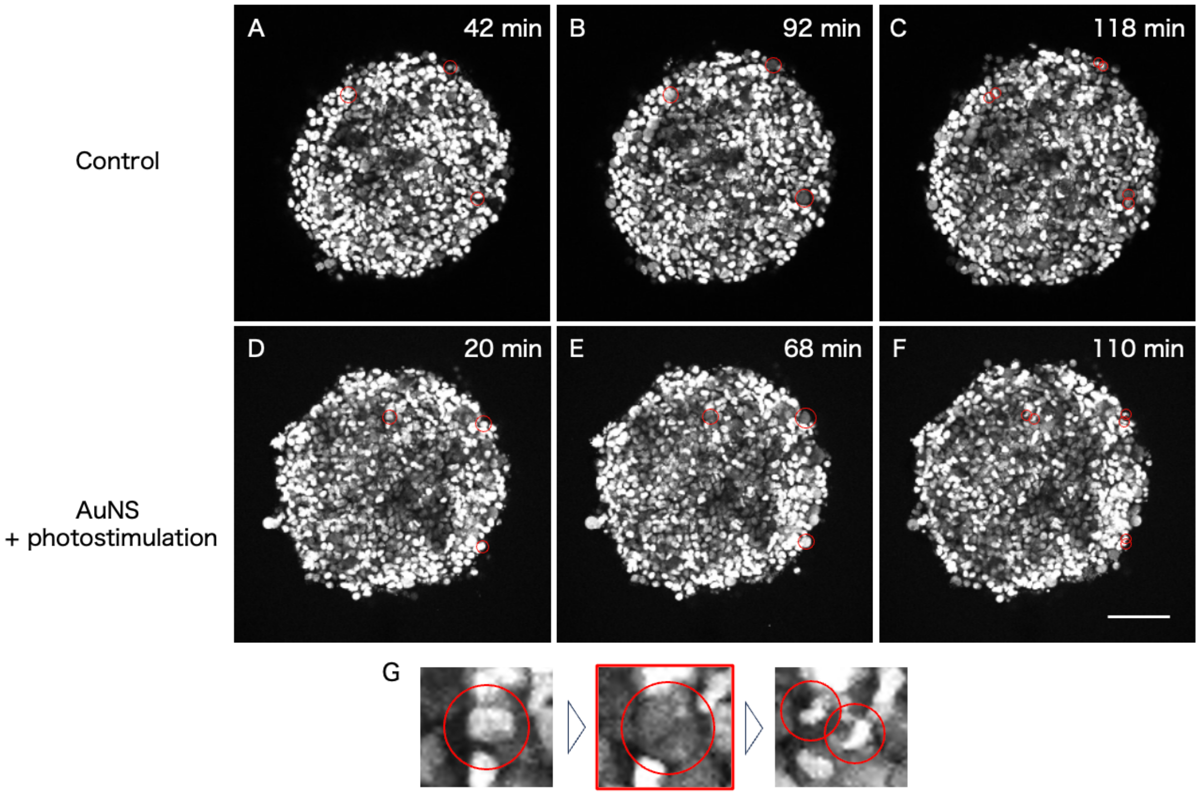


**Figure S6.** Maximum fluorescence intensity z projection of #T-HeLa spheroids (control) (A-C) and of #T-HeLa spheroids incubated with AuNSs (D-F) scanned at 1.5 mW mm$^{-2}$ (normal conditions). Red circles indicate points of cell division. The bottom insets show zoomed-in sections extracted from (D-F). Scale bar = 100 μm.
